# Producing Human Amniotic Epithelial Cells-only Membrane for Transplantation

**DOI:** 10.1101/837377

**Authors:** Chenze Xu, Waqas Ahmed, Lili Xie, Yan Peng, Qizheng Wang, Meijin Guo

## Abstract

Human amniotic epithelial cells (hAECs), as pluripotent stem cells, have characteristics of immune privilege and great clinical potential. Here, we produced hAECs membrane consisting of single-layer hAECs and basal membrane (BM) of human amniotic membrane (hAM). In conventional methods, hAECs were isolated from hAM by repeated trypsin digestion. In this study, collagenase I and cell scraper were used to remove human amniotic mesenchymal stem cells (hAMSCs) from hAM and hAECs-only membranes were produced. These hAECs on the membranes were evaluated by surface biomarkers including epithelial cell adhesion molecule (EpCAM), stage-specific embryonic antigen 4 (SSEA4) and endoglin (CD105), transcriptional level of biomarkers including POU class 5 homeobox 1 (OCT4), sex determining region Y-box 2 (SOX2), fibroblast growth factor 4 (FGF4), immunofluorescence of cytokeratin-8 (CK-8), alpha smooth muscle actin (α-SMA) and collagen type I alpha 1 chain (ColA1). Finally, the hAECs membrane were transplanted on skin-removed mice to evaluate its effect on wound healing. In comparison to the hAECs isolated by the conventional methods, the cells isolated by this proposed method had higher purity of hAECs, expressed higher in pluripotency related genes, and maintained an epithelium construction in a long-term culture. In mice application, the hAECs membrane effectively improved the skin wound healing. An efficient method was successfully established to produce hAECs membrane in this work which not only held promise to obtain hAECs in higher purity and quality, but also showed practical clinical potential.

## Introduction

hAECs are located on the surface of hAM by the side of umbilical cord. It’s proven that hAECs have a comprehensive pluripotency [1-3]. For instance, hAECs are capable of differentiation into osteoblasts [4], insulin-producing 3D spheroids [5], granulosa cells [6], corneal epithelial cells [7], hepatic cells [8], progenitor of cortical neurons [9], and pancreatic lineage [10]. hAECs have a characteristic of immune privilege due to a low expression of surface marker major histocompatibility complex, class II, DR (HLA-DR) [11]. Many studies have showed that hAECs have therapeutic potential on multiple sclerosis, immunomodulation, inflammation suppression, angiogenesis promotion, oxidative stress inhibition, neurogenesis induction, matrix metalloproteinases regulation, and remyelination stimulation [12-14]. Thus, hAECs can be applied to cellular transplantation for clinical use. However, there are still some obstacles in clinical application of hAECs.

One of the obstacles is that the purity of isolated hAECs still need improvement. In conventional methods, hAECs were isolated from hAM through multiple treatment of trypsin [15-21]. hAM are consist of three layers: single-layer of hAECs, a thick layer of BM which mainly made up of collagen and hyaluronic acid, and a thick layer of hAMSCs. Trypsin simultaneously affects the hAECs and hAMSCs. In most methods, the cumulative period of trypsin digestion was over 1 hour. Thus, many hAMSCs could be isolated together with hAECs which decreased the purity of hAECs. To address the issue, some researchers used hAECs and hAMSCs together in therapy [12,22]. However, hAECs and hAMSCs had different characteristics [4,23-25]. First, hAMSCs had differentiation potential of osteogenesis, chondrogenesis and adipogenesis, whereas hAECs had multidifferentiation potential. Thus, hAECs had much wider indications. Second, hAECs are epithelial cells and hAMSCs are mesenchymal cells. These two types of cells applied to different human tissue parts in transplantation. Although many researches had proved that hAMSCs could be capable of restricting the organ fibrosis and inhibiting the epithelial-to-mesenchymal transition, hAMSCs, as mesenchymal cells, could cause organ fibrosis by themselves in a long term [26-28]. Third, there are a lot of uncertain factors of the safety and reliability of hAMSCs in clinical application. For instance, in clinical trials, hAMSCs have risk of tumorigenicity [29,30], causing adverse responses like pain, swelling or heat [31-33], and even pneumonia-related death [34]. hAECs could also have some potential safety risks. Probably, it will increase much difficulty of safety risk evaluation in clinical trials when the test cells were mixed with hAECs and hAMSCs. Thus, it is significant to establish a novel approach to provide us with highly pure hAECs.

Another obstacle is that the hAECs should be made into epithelial sheet in epithelium transplantation therapy. To address the problem, some studies used several scaffolds for producing hAECs sheet including electrospun poly (lactide-co-glycolide), poly (ε-caprolactone), poly (lactic acid) scaffolds [35], and silk fibroin scaffold [36]. These techniques still faced some barriers to produce hAECs sheets of epithelium construction for reliable clinical application, including the cost of production, the immunogenicity of scaffold material, the cell adhesion capability of transplantation, the reduction degree of epithelium construction and etc. Thus, hAECs still need some other proper scaffolds for further therapeutic application.

To address the problems mentioned above, we produced hAECs membrane of epithelium construction by collagenase and cell scraper instead of trypsin. In previous studies, collagenase had been repeatedly tested, compared with trypsin, and turned out to be inefficient in isolation of hAECs [18-21]. One of the main reasons could be that collagenase affected both the BM and hAMSCs. Thus, a long term digestion by collagenase broked the BM into patches and released much collagen lysate which made it hard to separate hAECs and hAMSCs. In order to efficiently produce a sheet of highly pure hAECs membrane in this work, we first digested the hAM less than 30 min, kept the integrity of BM, and then scraped off the loose hAMSCs affected by collagenase I. Thus, a sheet of BM covered by hAECs (hAECs membrane) was produced. The hAECs membrane generated by this developed novel approach was proven to have a high purity and pluripotency of hAECs with an epithelium construction.

## Materials and methods

### Producing hAECs membrane

The Human placenta samples were aseptically processed in biosafety cabinet of class 10,000 clean room area. The hAM samples were isolated and cut into pieces of 5 cm, and weighed, performed as the established procedures [20]. The isolation methods of hAECs were tested including using tubes or plates, trypsin or collagenase I, different digestion period, and application of cell scrapers (**Supplementary Table S1, S2, S3**).

To produce hAECs membrane, some hAM samples were spread out in 9 cm plates with their hAECs side facing the bottom. The hAM samples were covered by 0.2% collagenase I reagent (Sigma, USA), followed by placing plates in rocking device (30-60 RPM) at 37°C for 5 min as pre-digestion. Then the samples were transferred into a new plate with fresh collagenase I reagent and incubated on rocking device (30-60 RPM) for less than 30 min (The exact processing period varied according to the actual digestion condition and digestion was terminated when hAMSCs began to peel off and the BM was undamaged). Collagenase I reagent was discarded and samples were washed with HBSS (Gibco, USA) for 2 times. Then cell scrapers were used on the hAMSCs side of the samples to remove the residual hAMSCs. followed by washing the samples with HBSS repeatedly until all the viscous liquid had been cleaned. Thus, the tissues left were hAECs membranes.

### hAECs Culture

The isolated hAECs and hAECs membrane were plated in 9 cm plates (1× 10^4^ cells/cm^2^), cultured in basic medium consisting of Dulbecco’s Modified Eagle Medium (DMEM) medium (low glucose, GlutaMAX, Pyruvate) (Gibco, USA) supplemented with 10% fetal calf serum (FBS) (Sigma, USA), 100 U/ml penicillin (Sigma, USA), 100 μg/ml streptomycin (Sigma, USA), 10ng/ml epidermal growth factor (EGF) (Gibco, USA), 25μM progesterone (Sigma, USA).

In mice experiment, three male mice were operated. All experimental operations were in accordance with laws, regulations and local ethical requirement in China.

### qPCR (Quantitative RT-PCR)

Total RNA from test groups was isolated using Invitrogen™ TRIzol™ (Thermo, USA), and reverse-transcribed by a PrimeScript™ RT reagent Kit with gDNA Eraser (Perfect Real Time) (TAKARA, Japan). qPCR was performed with SYBR Premix Ex Taq™ II (Tli RNaseH Plus) (TAKARA, Japan) according to the manufacturer’s instructions on a CFX96 touch qPCR system (Bio-Rad, USA). Primers used are listed in supplementary material (**Supplementary Table S6**)

### Immunofluorescence (IF) and Immunocytochemistry (ICC)

The cell samples being fixed with 4.0% paraformaldehyde solution (10-30min) were perforated on membrane by Triton X100 (0.1%, for less than 10min), and washed with phosphate buffer saline (PBS) for three times (10 min per wash). Later they were blocked with 5% bovine serum albumin (BSA) for 30 min, and incubated with antibodies and DAPI (Sigma, USA) according to manufactures’ instruction. Followed by washing with PBS as above, was incubated with secondary antibodies, before being completely ready for observation under an EVOS FL Auto imaging system (Life Technologies, USA). The antibodies used in this work are listed in supplementary material (**Supplementary Table S5**).

### Flow Cytometry (FCM) Analysis

The hAECs membrane was digested by 0.2% trypsin/EDTA (Thermo, USA) for 15 min at 37°C as suspended single cells, and then washed with PBS. For detecting intramembrane biomarkers, cell samples were treated with 4% paraformaldehyde solution for fixation (10-30 min) and then perforated on membrane by Triton X100 (0.1%, for about 10 min), later washed again with PBS (The procedures of cell membrane perforation were not performed in detecting cell surface markers). The samples were re-suspended in 100 μL volume of DMEM medium in a concentration of 1×10^6^-10^7^ cells/mL. Matched controls of antibodies for FCM were applied according to manufacturer’s instructions using a FACSArial system (BD Biosciences, USA). The Quad was set according to isotype antibodies. Main antibodies are listed in supplementary material (**Supplementary Table S4**).

### hAECs membrane transplantation on mice skin wound

Nine comatose 2-week-old C57BL6 black mice were prepared for surgical skin excision. First, these mice were shaved on the back and injected with 1% pelltobarbitalum patricum (10 mg/mL) for anesthetization according to each individual mouse (45 mg dose/kg weight). After confirmed unconscious, every mouse was surgically excised of a piece of square skin on their back in a thickness of 2 mm and a length of 1 cm. Then, six of them were transplanted by hAECs membrane on the skin wound and fixed by stitches. None of the mice died of bacterial infection or surgery.

Three of the transplanted mice were decapitated at 5 d. The wound skin samples were retrieved, sliced, and paraffined for immunohistochemistry (IHC). The rest were treated as above at 10 d.

### Statistical analysis

The hAM were derived from three donors. The investigation of isolation methods had 3 parallel samples in each group, respectively. A maximum and a minimum value were removed in each test group. In experiments of medium optimization and long-term culture, each test group was cultured on 9 cm plates with 5 parallel samples. In qPCR, cell number counting, and flow cytometry, results were the averages of 3 tests for each sample. To detect the expression of CK-8, α-SMA and Col1A1 antigens, the samples were treated with antibodies and Dapi (Invitrogen, USA), and counted of immune-positive results at 10 views per sample by immunofluorescence (IF). The complete procedures were successively repeated three times. Error bars represent ± SD. Reliable data meet the condition σ (Standard Deviation, SD) / μ (mean value) < 10%. Experimental data are reported as mean ± SD.

Statistical significance was evaluated by one-way ANOVA with SPSS software. *P*-value < 0.05 were considered statistically significant; *P*-value <0.01 have great significant statistical difference; *P*-value <0.001 have extreme great significant statistical difference.

## Results

### The cells on hAECs membrane results higher purity of hAECs

In conventional methods, hAECs were isolated through multiple digestion by trypsin [15-21,37]. The isolated cells by conventional methods could contain more hAMSCs and have lower purity of hAECs. Results of hematoxylin-eosin (HE) staining showed both of the hAECs side and hAMSCs side had obvious tissue defect (**Fig. 1F**). It proved that in general methods, the isolated cells were consist of both hAMSCs and hAECs. Here, the hAM were digested by collagenase I and scraped on the hAMSCs side. Results showed that most of the hAMSCs were removed from the hAM and the hAECs were unharmed (**Fig. 1F**). This tissue consisted of BM and single-layer of hAECs was identified as hAECs membrane (**Table 1**).

**Table 1.** The Methods of producing hAECs membrane and isolating hAECs.

| **Procedures for producing hAECs membrane** |  | **Viabi**<br>**-lity** | **Cell number**<br>**(cells/g)** | **EpCAM<sup>+</sup>**<br>**cells** |
| --- | --- | --- | --- | --- |
| Step 1 | The samples were placed on plates with the hAMSCs side up, digested with 0.2% Collagenase I for 5 min as pre-digestion, and then for less than 30 min. | 88.4% | 3.21±0.09<br>×10<sup>6</sup> | 2.32% |
| Step 2 | The samples were scraped on the hAMSCs side. Washed twice with PBS and retrieve all the solution for cell collection. | 85.6% | 1.97±0.39<br>×10<sup>6</sup> | 6.54% |
| Step 3 | The tissue left was hAECs membrane. hAECs membrane was digested by trypsin for 15 min and the cells were collected for characterization. | 79.1% | 1.83±0.21<br>×10<sup>6</sup> | 92.2% |
| **Procedures of hAECs isolation by trypsin** |  | **Viabi**<br>**-lity** | **Cell number**<br>**(cells/g)** | **EpCAM<sup>+</sup>**<br>**cells** |
| Step 1 | The samples were placed on plate with hAECs side up, digested with 0.2% Trypsin/EDTA for 15 min as pre-digestion, and then for 30 min. Collect cells in the digestion solution. | 71.4% | 2.21±0.31<br>×10<sup>6</sup> | 86.1% |
| Step 2 | The samples were digested in 0.2% Trypsin/EDTA in tubes for 30 min. | 61.8% | 3.65±0.46<br>×10<sup>6</sup> | 32.7% |

**Fig. 1.**
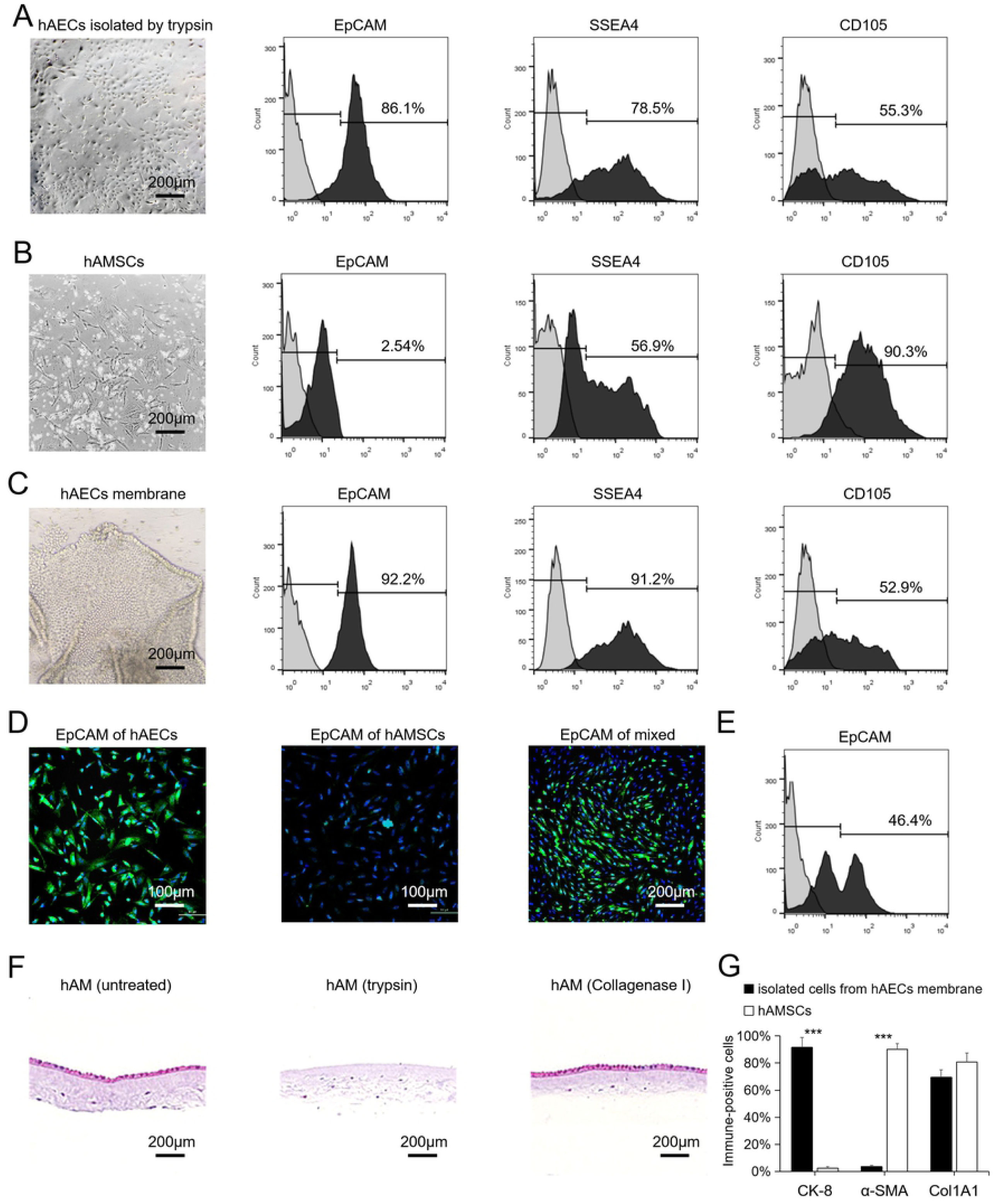
Characterize the isolated cells from hAM. (A) hAECs isolated by trypsin through the existed methods [20]. The immune-positive result of EpCAM, SSEA4 and CD105 was set according to the result of antibody isoform, hAECs isolated by trypsin and hAMSCs. Scale bar = 200μm. (B) hAMSCs isolated by the existed methods [40]. (C) Shows produced hAECs membrane. The cells digested by trypsin as single cells for flow cytometry. (D) hAECs isolated by trypsin, hAMSCs and the mixed of the two cells identified of EpCAM antigen through IF. hAECs showed EpCAM^+^. hAMSCs showed EpCAM^−^. The mixed cells showed distinct expression of EpCAM. (E) The mixed cells of hAECs and hAMSCs showed two separated peaks indicating EpCAM with different expression between hAECs and hAMSCs. (F) Left untreated hAM. Middle the hAM treated by trypsin according to existed methods [20]. Right the hAM treated by Collagenase I and cell scraper. (G) The immune-positive results of CK-8, α-SMA, and Col1A1 detected in hAECs from hAECs membrane and hAMSCs through IF. The immune- positive results were counted in ten view each sample. Results were expressed as mean ± SD (n = 3 independent experiments). Asterisks indicate statistical significance of differences in the immune-positive results between the hAECs and hAMSCs. (*-P-value <0.05, **-P-value <0.01, ***-P-value <0.001).

To demonstrate that the hAECs membrane comprised pure hAECs, the isolated cells from different methods were identified by some specific biomarkers. To distinctly identify hAECs and hAMSCs, the most accepted biomarkers to distinguish hAECs and hAMSCs were EpCAM (hAECs-positive), CK-8 and CK-18 (hAECs-positive) and α-SMA (hAECs-negative). Results showed that hAMSCs isolated by former established methods had 2.54% EpCAM^+^ result [4,23,38] (**Fig. 1B**) (**Table 1**). The hAECs isolated by general methods had 86.1% positive result of EpCAM. According to the samples mixed with hAECs and hAMSCs (from Method B, Step 2a), the samples had two clear separated peaks by flow cytometry and showed 46.4% EpCAM^+^ result (**Fig. 1E**). In IF analysis, the hAMSCs isolated by established procedures showed EpCAM^−^ [4,23,38] (**Fig. 1D**). The hAECs derived from hAECs membrane showed EpCAM^+^. The samples of mixed hAECs and hAMSCs (from Method B, Step 2a) had obvious difference between positive or negative expression of EpCAM. Thus, it’s proven that hAMSCs expressed low in EpCAM antigen while hAECs had high expression. These evidences indicate that EpCAM can be used as the main specific biomarker of hAECs. According to the EpCAM biomarker, the purity of hAECs on hAECs membrane (92.2%) was proven to be higher than those isolated by conventional methods (86.1%). Furthermore, hAMSCs had relatively low immune-positive result of SSEA4 (56.9%). The cells from hAECs membrane (91.2%) showed higher SSEA4^+^ than those isolated by conventional methods (78.5%). Thus, it also implied that the cells on hAECs membrane had higher purity of hAECs.

For further identification, the cells were compared on CK-8, α-SMA and Col1A1 markers through IF analysis. Results showed that the isolated cells from hAECs membrane expressed high in CK-8 and low in α-SMA, and similar in Col1A1 compare to hAMSCs (**Fig. 1G**). These results were according with the former findings in characteristic of hAECs [9,16,38-40]. Therefore, it showed that the cells on hAECs membrane generated here were very likely to be highly pure hAECs.

Regarding the viability of the cells isolated from hAECs membrane, the viability of the isolated cells by general methods was 71.4% (recovered from first digestion) and 61.8% (recovered from second digestion) (**Table 1**). In comparison, the viability of the cells isolated from hAECs membrane produced here was 79.1%. Thus, the cells on hAECs membrane had higher viability.

The hAECs membrane was produced through removing the hAMSCs by combination of collagenase I and cell scraper. Results indicate that the cells from hAECs membrane show higher purity of hAECs, better pluripotency and higher viability compare to those isolated by methods previously described. Conclusively, it was an alternative efficient approach of isolating highly pure hAECs.

### The hAECs membrane exhibited epithelial construction and performed better in characteristic maintenance

In general methods, hAECs were isolated into single cells and cultured. In this study, the hAECs were cultured as one-layer on BM which had potential to mimic an *in vivo* micro-environment for these hAECs.

To identify the difference between the hAECs cultured as single cells and those on hAECs membrane, two test groups were compared in specific biomarker, pluripotency related biomarkers and genes. In group (hAECs membrane), a hAECs membrane was spread out on a 9 cm plate with BM side facing the bottom and cultured in SCM. In group (hAECs separated), the single cells isolated from hAECs membranes by trypsin were cultured in SCM (2 × 10^4^ cells/cm^2^). After 15 d, the cells of cultured hAECs membrane orderly arranged with similar cellular size and clear edge (**Fig. 2B**). On the contrary, the separated hAECs grew into patches in different cellular size, irregular edge shape, and excessively attached to each other (**Fig. 2A**). Thus, the results indicated that the hAECs membrane had a more similar construction of epithelium than those hAECs cultured as single cells.

**Fig. 2.**
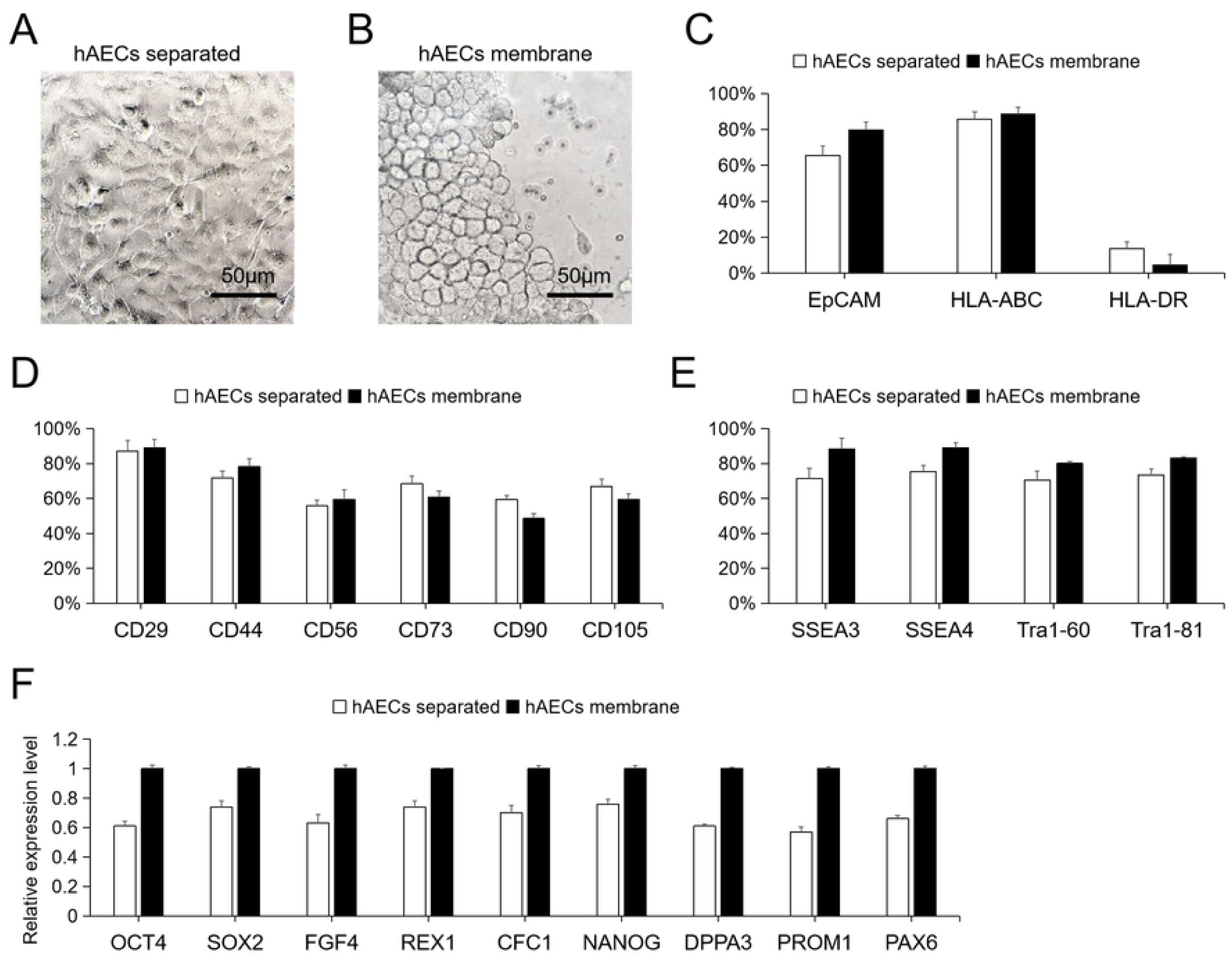
Difference between the separated hAECs and the cells on hAECs membrane in long-term culture. (A) The hAECs separated from hAECs membrane cultured for 15 d. (B) The hAECs membrane planted on a plate and cultured for 15 d. The hAECs separated and those on hAECs membrane were compared in (C) specific surface marker EpCAM and immunity related surface markers, (D) other specific surface markers, and (E) pluripotency related surface markers. (F) The transcripition expression results of pluripotency related genes expressed relative to the higher mean value between the hAECs separated and those on hAECs membrane. Results showed as mean ± SD (n = 3 independent experiments).

Afterwards, the cells of group (hAECs separated) and group (hAECs membrane) were determined through flow cytometry and qPCR. Results showed the cells on the hAECs membrane had more immune-positive specific biomarker, EpCAM; MHC I marker, HLA-ABC; other biomarkers including CD29, CD44, CD56; and much less immune-positive results of MHC II maker HLA-DR (**Fig. 2C, 2D**). These results indicated that the cells on hAECs membrane had better maintenance of their own main characteristic biomarkers. Human immune system mainly recognizes the allogeneic cells through MHC II surface antigen. The immune-positive result of HLA-DR, as a MHC II antigen, was quite low in hAECs membrane cultured for 15 d (**Fig. 2C**). It indicated that the cells on hAECs membrane had low immunogenicity and maintained to be applicable for tissue transplantation after a long-term culture (**Fig. 2C**). The transcriptional level of pluripotency related genes OCT4, SOX2, FGF4, REX1, CFC1, NANOG, DPPA3, PROM1 and PAX6, and immune-positive results of pluripotency related biomarkers SSEA3, SSEA4, Tra1-60, Tra1-80 and GCTM2 were higher in hAECs membrane than in separated hAECs (**Fig. 2E, 2F**). These results showed that the hAECs membrane had prolong the maintenance of the pluripotency of the hAECs.

Based on the results, the cells on the hAECs membrane had a similar construction of epithelium and better maintenance of pluripotency, immune privilege, and specific characteristic compare to those hAECs cultured as single cells. It was speculated that the hAECs membrane had provided the hAECs with some functional factors or a more appropriate micro-environment.

### Transplantation of hAECs membrane for healing mice skin wound

It had been proven that the BM of hAM had great therapeutic potential in human ocular surface reconstruction [41-43], and wound healing caused by radiation [44]. Also, hAECs were used in tissue regeneration for their characteristic of pluripotency and immune privilege [12,13,45]. Thus, the hAECs membrane could be potential medical treatment for skin and endothelium wound, psoriasis, lupus erythematosus, diabetes-caused ulceration, etc. In this study, the hAECs membrane was used to heal mice skin wound.

Nine black mice were surgically excised of skin surface in thickness of 2 mm, and six of them were transplanted of hAECs membrane. After 5 d, the hAECs membrane closely attached on the wound area and prevented the dermis from exposure to the air (**Fig. 3B**). At 10 d, the skin wound of control group was heavily scabby (**Fig. 3D**). In transplanted mice, no sign of scab, scar formation, or exposed dermis were observed (**Fig. 3C**). In the tissue slider of mice skin wound area, the transplanted hAECs membrane had conjugated to the dermis at 5 d (**Fig. 3F**). At 10 d, there were corneum-like construction formed around hAECs (**Fig. 3G**). The BM was degraded and fell off. In comparison, the surface of skin wound of mice in control group were covered by thick scab, and there was no apparent regeneration of dermis, epithelium and corneum (**Fig. 3H**). Thus, results showed that the skin wound performed better regeneration by the transplantation of hAECs membrane.

**Fig. 3.**
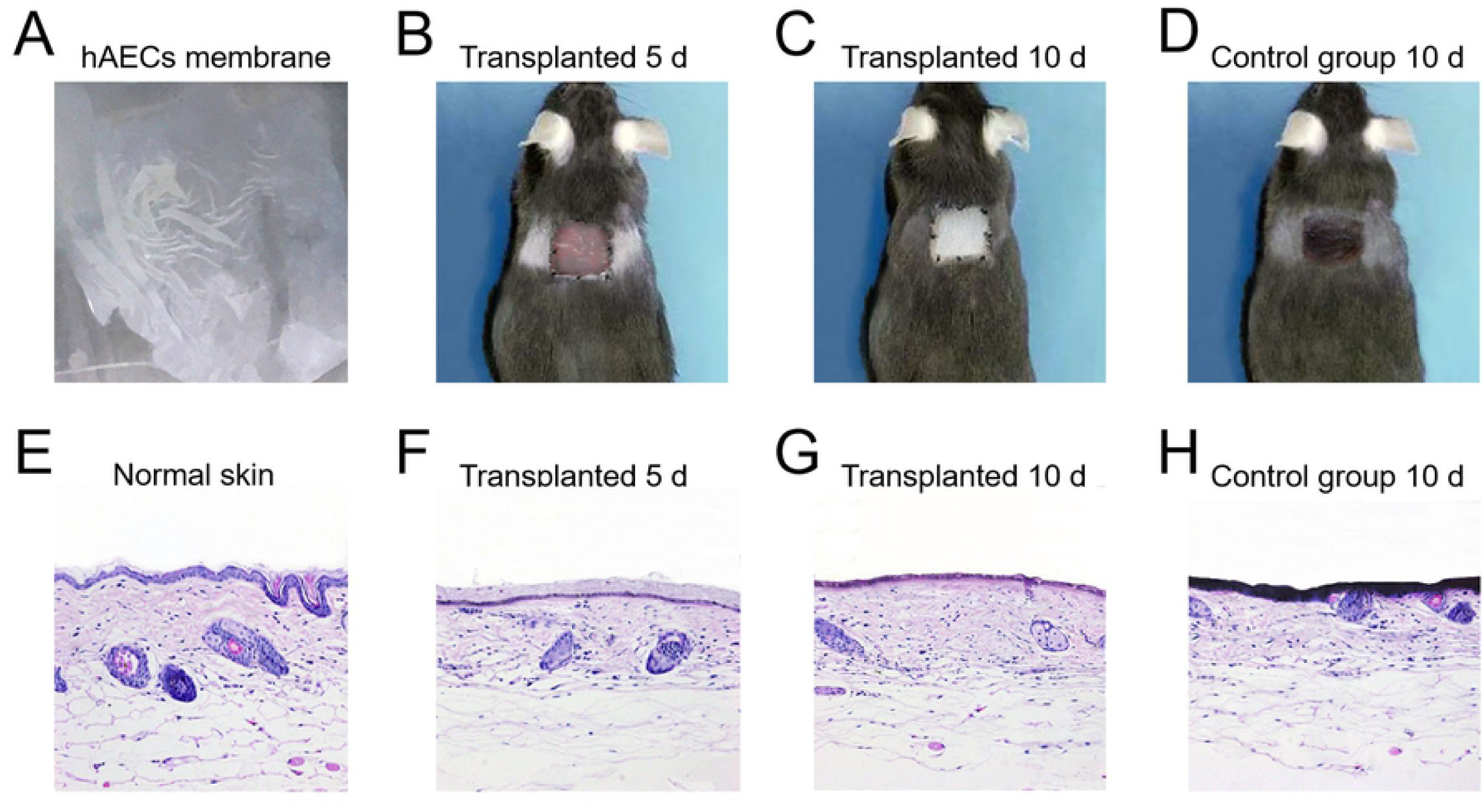
Mice skin wound healing test. (A) The hAECs membrane showed as a white, thin translucent membrane. (B) hAECs membrane had been transplanted on the skin wound at 5 d. There was no sign of bleeding, inflammation or further aggravation. (C) Skin wound had been repaired by hAECs membrane at 10 d. There was no sign of scar formation or inflammation. (D) Skin wound had thick scab in mouse of control group at 10 d. (E) A slide of normal skin. (F) At 5 d, the slide showed hAECs membrane had been conjugated to the mouse dermis. (G) At 10 d, the slide showed the skin surface had epithelium-like and cornium-like reconstruction. (H) At 10 d, the slide showed a thick scab on the skin wound.

## Discussion

### Key points to efficiently produce hAECs membrane

In recent years, many studies had established methods of isolating hAECs [15-21]. However, the characteristic of these hAECs from different methods were quite different. One of the main reason was these methods could have caused uneven efficiency and purity of hAECs. Thus, we proposed a novel alternative approach to isolate highly pure hAECs by removing hAMSCs from hAM with collagenase I and cell scraper. There were several key points influencing the production of hAECs membrane.

First, based on the tested methods, it was important to use plates instead of tubes to remove the hAMSCs. The cells isolated by collagenase I were more in number and lower in EpCAM^+^ result on plates (Method F, Step 1, 2.32% of EpCAM^+^ in 3.21±0.09×10^6^ cells/g hAM) than those in tubes (Method B, Step 1, 4.52% of EpCAM^+^ in 4.67± 0.53×10^5^ cells/g hAM) (**Supplementary Table S1, S3**). Thus, the hAMSCs were more efficiently and selectively removed when the digestion was performed on the plates. One of the main reasons was that the hAM samples had an uneven digestion in tubes because of discrepant exposure of collagenase. On the contrary, the hAM samples on the plates stretched, were evenly digested, and released less collagen lysate.

Second, it was necessary to place the hAM samples with their hAMSCs side facing up. Results showed the EpCAM^−^ cells were more efficiently isolated in Method F, Step 1 when the hAMSCs side faced up (2.32% of EpCAM^+^ in 3.21±0.09×10^6^ cells/g hAM) than in Method E, Step 1 when the hAECs side faced up (1.84% of EpCAM^+^ in 4.86±0.25×10^4^ cells/g hAM) (**Supplementary Table S3**). These results indicated the hAMSCs were supposed to face up to be efficiently removed. Probably, the hAMSCs side faced up, were more likely to be exposed to fresh collagenase I and avoided the collagen lysate derived from BM and hAMSCs.

Third, the repeated treatment with combination of collagenase digestion and cell scraping improved the efficiency and success rate in producing hAECs membrane. In digestion, collagen lysate released, filled the gap between cells, and constructed the digestion process. Thus, cell scraper was necessary to remove the loosened hAMSCs and collagen lysate from the surface. In practical operation, regular scraping the surface of hAMSCs side of these hAM samples can improve the efficiency of removing hAMSCs and prevent the excessive digestion of BM.

Fourth, different digestion condition affected the thickness of BM of the hAECs membrane. The digestion period and volume of Collagenase I depended on the inter-individual difference of placenta donors, thickness and quality of hAM samples, and the quantity of released collagen lysate. If the digestion was insufficient, there were much residue hAMSCs attached on the hAECs membrane. When the digestion was excessive, the hAECs membrane were damaged or broken. Thus, the hAM samples need to be observed under microscope every 5 min to evaluate the digestion degree.

Finally, the residue of red blood cells should be washed off as much as possible. In culture, it was observed that the red blood cells obstructed the attachment and migration of the hAECs or hAMSCs. It was a critical point to clean off the blood at the time of sample collection.

### Characterization of the hAECs membrane

In this study, a hAECs membrane was produced by collagenase I and cell scraper. Based on the results, the cells from the hAECs membrane showed higher viability, purity, and better pluripotency than the hAECs isolated by the general methods.

In conventional methods, the hAM samples were repeatedly treated by trypsin and the whole digestion period reached 75 min. Based on the results, the viability of the isolated cells treated by trypsin was 71.4% for 45 min, was for 61.8% for 75 min (**Table 1**). In comparison, the hAECs isolated by collagenase I in 35 min and trypsin for 15 min was 79.1%. Thus, the isolated hAECs had much higher viability by the methods established in this study. There were two main reasons: 1. Collagenase I had little influence on the hAECs, but worked through digesting the collagen on the surface structure of hAMSCs [38,46,47]. Thus, the application of Collagenase I did not cause much damage to the hAECs; 2. The digestion period in producing hAECs membrane was much shorter than the conventional methods. In conventional methods, the hAM was treated by trypsin for 15 min, 30 min, and then another 30 min. The hAECs were considered to suffer a certain damage on their cell membrane or even disrupted. In the approach to produce hAECs membrane, the samples merely suffered from 0.2% collagenase I for 30 min. Yet, these hAECs were still unharmed. The viability of the isolated cells did not just indicate the living cells’ portion, but also used to identify the global state of these cells. In the isolated cells with low viability, these living cells could have suffered different levels of damage which caused negative results in proliferation, survival, pluripotency and differentiation capability. To accord with this view, results showed that the hAECs from hAECs membrane had higher pluripotency marker SSEA4^+^ result (91.2%) than those by general methods (78.5%).

In addition, results indicated that the hAECs membrane had higher purity of hAECs (92.2% EpCAM^+^) than those by general methods (86.1% EpCAM^+^). In analysis, the high purity was resulted from the selectivity of collagenase. In general methods, trypsin worked on both the hAECs and hAMSCs. Thus, the application of trypsin was efficient to selectively separate hAECs and hAMSCs. In producing hAECs membrane, Collagenase I selectively digested the collagen in the surface structure of hAMSCs. With proper scraping, most of hAMSCs were removed from hAM, and only hAECs attached on the BM. Therefore, the procedure of producing hAECs membrane was more selective in isolating hAECs. However, the EpCAM^+^ result of the hAECs membrane was not up to 95%. There could be several reasons: 1. Not all of these hAECs expressed high in EpCAM surface marker antigen; 2. There were residue hAMSCs left on the hAM. 3. Some hAMSCs were adhered on the hAECs membrane through the viscous collagen lysate. Thus, it remains a task to improve the methods of producing hAECs membrane for raising the purity and cell states of hAECs.

### The hAECs membrane could provide a better micro-environment for hAECs’ culture

Generally, hAECs isolated from hAM were not able to maintain their characteristic for a long time. In 2 weeks, hAECs gradually lost the expression of specific markers like EpCAM, CD29, and CD44; pluripotency markers like SSEA3 and SSEA4; and pluripotency related genes like OCT4 and SOX2 [15,16,24,39]. Then, these hAECs could happen epithelial-to-mesenchymal transition, or spontaneous differentiation, which were adverse to their clinical application. To address these problems, some studies used different factors such as progesterone, EGF or Supplement S7in hAECs’ culture [5,15-19,21,39,48]. However, these results were still not ideal.

In this study, it was found that hAECs can maintain in a good condition when cultured as hAECs membrane. The immune-positive results of some specific markers EpCAM, CD29, CD44, and CD56, pluripotency markers SSEA3, SSEA4, Tra1-60, and Tra1-80 had maintained in a high level (**Fig. 2C, 2D, 2E**). In addition, the transcription level of pluripotency related genes OCT4, SOX2, FGF4, REX1, CFC1, NANOG, DPPA3, PROM1, and PAX6 was much higher in the cells of hAECs membrane (**Fig. 2F**). Accordingly, the hAECs membrane was speculated to provide a better micro-environment for the maintainance of hAECs. There could be three main reasons. First, the hAECs membrane immobilized the hAECs and kept them in an epithelium construction, exactly as how they grow *in vivo*. In general methods, hAECs were cultured as single cells. Based on the results, the hAECs separated as single cells disorderly grew into single-layer (**Fig. 2A**). In the meanwhile, the cells on hAECs membrane still had an epithelium construction (**Fig. 2B**). Thus, these hAECs could have a proper surface contact on the hAECs membrane.

Second, the BM of the hAECs membrane could have provided a better support surface. Generally, mammalian cells have a high demand of a proper support surface. Thus, some material such as gelatin, collagen matrix and matrigel (BD, USA) were used as support surface for culture of pluripotent stem cells like embryonic stem cells or induced pluripotent stem cells[49-52]. The BM of hAM was mainly consisted of collagen and proven to be a good supporting scaffold [12,53]. Thus, it was speculated that the BM could properly support the hAECs in culture. Furthermore, the hAECs were conjugated to BM *in vivo*. Thus, they could have high compatibility.

Third, the BM of hAECs membrane could reserve some factors helping the maintenance of hAECs. In general methods, hAECs were isolated as single cells and most of the protein factors had been washed off. In this approach of producing hAECs membrane, some protein factors could still be stored in BM, and brought positive influence on hAECs for a long time.

Thus, the hAECs membrane was speculated to provide good micro-environment for hAECs’ culture by preserving epithelium construction, providing proper support surface and valuable protein factors.

## Conclusion

A novel alternative approach of producing hAECs membrane was established in this study. This hAECs membrane had high purity of hAECs and epithelium construction, improved maintenance of pluripotency and immune privilege, and were capable of improving the skin wound healing on mice, which showed great potential for the future clinical application.

## Declarations

The collection process of placenta from voluntary donors were approved by the ethics committee led by Minxiang Lao, the director of Biological clinic center of the Shanghai Renji Hospital, through signed informed consent from the mothers tested for human immunodeficiency virus, Hepatitis B virus, C virus, tuberculosis, *Chlamydia trachomatis, Neisseria gonorrhoeae* and Syphilis, and the healthy mothers underwent elective cesarean deliveries. The ethics committee included Doctor Xia Wu, Jie Zhu, Yunyan Chen, Yi Jin and Fang Ji.

The laboratory mice were acquired from Shanghai Jie Si Jie laboratory animal (JSJ-lab) company with Permit for use of laboratory animals (SYXK) (China), and approved for research purpose by the local ethics committee. Nine male mice had operations according to European animal care guidelines. All experimental operations were in accordance with laws, regulations and local ethical requirement in China.

## Consent for publication

Not applicable.

## Availability of data and material

The data supporting the findings was list in Supplementary Material uploaded with the manuscript.

## Acknowledgments

We greatly appreciate the donor for placenta samples and the hard work of all the participants.

## Author contributions

This study was designed by MG & CX and most experiments were performed by CX with help from LX, YP and QW. The experimental data was also analyzed by CX. The paper was written by CX, WA and MG. MG and WA participated in the data discussion.

## Reference

1. Miki T, T Lehmann, H Cai, DB Stolz and SC Strom. (2005). Stem cell characteristics of amniotic epithelial cells. Stem Cells 23:1549–59.

2. Fanti M, R Gramignoli, M Serra, E Cadoni, SC Strom and F Marongiu. (2017). Differentiation of amniotic epithelial cells into various liver cell types and potential therapeutic applications. Placenta 59:139–145.

3. Miki T. (2018). Stem cell characteristics and the therapeutic potential of amniotic epithelial cells. Am J Reprod Immunol 80:e13003.

4. Shen C, C Yang, S Xu and H Zhao. (2019). Comparison of osteogenic differentiation capacity in mesenchymal stem cells derived from human amniotic membrane (AM), umbilical cord (UC), chorionic membrane (CM), and decidua (DC). Cell Biosci 9:17.

5. Okere B, F Alviano, R Costa, D Quaglino, F Ricci, M Dominici, P Paolucci, L Bonsi and L Iughetti. (2015). In vitro differentiation of human amniotic epithelial cells into insulin-producing 3D spheroids. Int J Immunopathol Pharmacol 28:390–402.

6. Tuan RS and T O’Brien. (2015). Expression of Concern: Human amniotic epithelial cells can differentiate into granulosa cells and restore folliculogenesis in a mouse model of chemotherapy-induced premature ovarian failure. Stem Cell Res Ther 6:240.

7. Liu X, J Chen, Q Zhou, W Xu, L Ye, J Zhou and L Jin. (2017). [Differentiation of human amniotic epithelial cells (HAECs) into corneal epithelial cells induced by co-culture of HAECs and corneal epithelial cells in vitro]. Xi Bao Yu Fen Zi Mian Yi Xue Za Zhi 33:508–512.

8. Maymo JL, R Riedel, A Perez-Perez, M Magatti, B Maskin, JL Duenas, O Parolini, V Sanchez-Margalet and CL Varone. (2018). Proliferation and survival of human amniotic epithelial cells during their hepatic differentiation. PLoS One 13:e0191489.

9. Garcia-Castro IL, G Garcia-Lopez, D Avila-Gonzalez, H Flores-Herrera, A Molina-Hernandez, W Portillo, E Ramon-Gallegos and NF Diaz. (2015). Markers of Pluripotency in Human Amniotic Epithelial Cells and Their Differentiation to Progenitor of Cortical Neurons. PLoS One 10:e0146082.

10. Balaji S, Y Zhou, EC Opara and S Soker. (2017). Combinations of Activin A or Nicotinamide with the Pancreatic Transcription Factor PDX1 Support Differentiation of Human Amnion Epithelial Cells Toward a Pancreatic Lineage. Cell Reprogram 19:255–262.

11. Yang PJ, WX Yuan, J Liu, JY Li, B Tan, C Qiu, XL Zhu, C Qiu, DM Lai, LH Guo and LY Yu. (2018). Biological characterization of human amniotic epithelial cells in a serum-free system and their safety evaluation. Acta Pharmacol Sin 39:1305–1316.

12. Farhadihosseinabadi B, M Farahani, T Tayebi, A Jafari, F Biniazan, K Modaresifar, H Moravvej, S Bahrami, H Redl, L Tayebi and H Niknejad. (2018). Amniotic membrane and its epithelial and mesenchymal stem cells as an appropriate source for skin tissue engineering and regenerative medicine. Artif Cells Nanomed Biotechnol 46:431–440.

13. Muttini A, B Barboni, L Valbonetti, V Russo and N Maffulli. (2018). Amniotic Epithelial Stem Cells: Salient Features and Possible Therapeutic Role. Sports Med Arthrosc Rev 26:70–74.

14. Abbasi-Kangevari M, SH Ghamari, F Safaeinejad, S Bahrami and H Niknejad. (2019). Potential Therapeutic Features of Human Amniotic Mesenchymal Stem Cells in Multiple Sclerosis: Immunomodulation, Inflammation Suppression, Angiogenesis Promotion, Oxidative Stress Inhibition, Neurogenesis Induction, MMPs Regulation, and Remyelination Stimulation. Front Immunol 10:238.

15. Tabatabaei M, N Mosaffa, S Nikoo, M Bozorgmehr, R Ghods, S Kazemnejad, S Rezania, B Keshavarzi, S Arefi, F Ramezani-Tehrani, E Mirzadegan and AH Zarnani. (2014). Isolation and partial characterization of human amniotic epithelial cells: the effect of trypsin. Avicenna J Med Biotechnol 6:10–20.

16. Liu X, Q Zhou, X Zhang, X Qin, W Wang, L Wang, J Meng and J Chen. (2014). [In vitro culture and molecular characterization of human amniotic epithelial cells]. Xi Bao Yu Fen Zi Mian Yi Xue Za Zhi 30:1318–21.

17. Murphy SV, A Kidyoor, T Reid, A Atala, EM Wallace and R Lim. (2014). Isolation, cryopreservation and culture of human amnion epithelial cells for clinical applications. J Vis Exp.

18. Lai D, Y Wang, J Sun, Y Chen, T Li, Y Wu, L Guo and C Wei. (2015). Derivation and characterization of human embryonic stem cells on human amnion epithelial cells. Sci Rep 5:10014.

19. Gramignoli R, RC Srinivasan, K Kannisto and SC Strom. (2016). Isolation of Human Amnion Epithelial Cells According to Current Good Manufacturing Procedures. Curr Protoc Stem Cell Biol 37:1e.10.1-1e.10.13.

20. Motedayyen H, N Esmaeil, N Tajik, F Khadem, S Ghotloo, B Khani and A Rezaei. (2017). Method and key points for isolation of human amniotic epithelial cells with high yield, viability and purity. BMC Res Notes 10:552.

21. Gottipamula S and KN Sridhar. (2018). Large-scale Isolation, Expansion and Characterization of Human Amniotic Epithelial Cells. Int J Stem Cells 11:87–95.

22. Yu SC, YY Xu, Y Li, B Xu, Q Sun, F Li and XG Zhang. (2015). Construction of tissue engineered skin with human amniotic mesenchymal stem cells and human amniotic epithelial cells. Eur Rev Med Pharmacol Sci 19:4627–35.

23. Gan WT, X Sun and Y Lu. (2015). [Comparison of Biological Characteristics between Human Amnion Epithelial Cells and Human Amnion Mesenchymal Stem Cells]. Zhongguo Shi Yan Xue Ye Xue Za Zhi 23:1120–4.

24. Wu Q, T Fang, H Lang, M Chen, P Shi, X Pang and G Qi. (2017). Comparison of the proliferation, migration and angiogenic properties of human amniotic epithelial and mesenchymal stem cells and their effects on endothelial cells. Int J Mol Med 39:918–926.

25. Ran LJ, Y Zeng, SC Wang, DS Zhang, M Hong, SY Li, J Dong and MX Shi. (2018). Effect of coculture with amniotic epithelial cells on the biological characteristics of amniotic mesenchymal stem cells. Mol Med Rep 18:723–732.

26. Purdon S, CL Patete and MK Glassberg. (2019). Multipotent Mesenchymal Stromal Cells for Pulmonary Fibrosis? Am J Med Sci 357:390–393.

27. Rong X, J Liu, X Yao, T Jiang, Y Wang and F Xie. (2019). Human bone marrow mesenchymal stem cells-derived exosomes alleviate liver fibrosis through the Wnt/beta-catenin pathway. Stem Cell Res Ther 10:98.

28. Zhang Y, X Jiang and L Ren. (2019). Optimization of the adipose-derived mesenchymal stem cell delivery time for radiation-induced lung fibrosis treatment in rats. Sci Rep 9:5589.

29. Barkholt L, E Flory, V Jekerle, S Lucas-Samuel, P Ahnert, L Bisset, D Buscher, W Fibbe, A Foussat, M Kwa, O Lantz, R Maciulaitis, T Palomaki, CK Schneider, L Sensebe, G Tachdjian, K Tarte, L Tosca and P Salmikangas. (2013). Risk of tumorigenicity in mesenchymal stromal cell-based therapies--bridging scientific observations and regulatory viewpoints. Cytotherapy 15:753–9.

30. Casiraghi F, G Remuzzi, M Abbate and N Perico. (2013). Multipotent mesenchymal stromal cell therapy and risk of malignancies. Stem Cell Rev 9:65–79.

31. Kang MH and HM Park. (2014). Evaluation of adverse reactions in dogs following intravenous mesenchymal stem cell transplantation. Acta Vet Scand 56:16.

32. Joswig AJ, A Mitchell, KJ Cummings, GJ Levine, CA Gregory, R Smith, 3rd and AE Watts. (2017). Repeated intra-articular injection of allogeneic mesenchymal stem cells causes an adverse response compared to autologous cells in the equine model. Stem Cell Res Ther 8:42.

33. Ursini TL, LL Amelse, HA Elkhenany, A Odoi, JL Carter-Arnold, HS Adair and MS Dhar. (2019). Retrospective analysis of local injection site adverse reactions associated with 230 allogenic administrations of bone marrow-derived mesenchymal stem cells in 164 horses. Equine Vet J 51:198–205.

34. Forslow U, O Blennow, K LeBlanc, O Ringden, B Gustafsson, J Mattsson and M Remberger. (2012). Treatment with mesenchymal stromal cells is a risk factor for pneumonia-related death after allogeneic hematopoietic stem cell transplantation. Eur J Haematol 89:220–7.

35. Russo V, L Tammaro, L Di Marcantonio, A Sorrentino, M Ancora, L Valbonetti, M Turriani, A Martelli, C Camma and B Barboni. (2016). Amniotic epithelial stem cell biocompatibility for electrospun poly(lactide-co-glycolide), poly(epsilon-caprolactone), poly(lactic acid) scaffolds. Mater Sci Eng C Mater Biol Appl 69:321–9.

36. Wang TG, J Xu, AH Zhu, H Lu, ZN Miao, P Zhao, GZ Hui and WJ Wu. (2016). Human amniotic epithelial cells combined with silk fibroin scaffold in the repair of spinal cord injury. Neural Regen Res 11:1670–1677.

37. Topoluk N, R Hawkins, J Tokish and J Mercuri. (2017). Amniotic Mesenchymal Stromal Cells Exhibit Preferential Osteogenic and Chondrogenic Differentiation and Enhanced Matrix Production Compared With Adipose Mesenchymal Stromal Cells. Am J Sports Med 45:2637–2646.

38. Deng Y, G Huang, L Zou, T Nong, X Yang, J Cui, Y Wei, S Yang and D Shi. (2018). Isolation and characterization of buffalo (bubalus bubalis) amniotic mesenchymal stem cells derived from amnion from the first trimester pregnancy. J Vet Med Sci 80:710–719.

39. Pratama G, V Vaghjiani, JY Tee, YH Liu, J Chan, C Tan, P Murthi, C Gargett and U Manuelpillai. (2011). Changes in culture expanded human amniotic epithelial cells: implications for potential therapeutic applications. PLoS One 6:e26136.

40. Barbati A, M Grazia Mameli, A Sidoni and GC Di Renzo. (2012). Amniotic membrane: separation of amniotic mesoderm from amniotic epithelium and isolation of their respective mesenchymal stromal and epithelial cells. Curr Protoc Stem Cell Biol Chapter 1:Unit 1E.8.

41. Bhatia J, B Narayanadas, M Varghese, MA Mansi, MF Mohamed, AAH Gad and N Bhatia. (2019). Management of recalcitrant corneal ulcers with dry processed amniotic membrane. Oman J Ophthalmol 12:68–69.

42. Bandeira F, GH Yam, M Fuest, HS Ong, YC Liu, XY Seah, SY Shen and JS Mehta. (2019). Urea-De-Epithelialized Human Amniotic Membrane for Ocular Surface Reconstruction. Stem Cells Transl Med.

43. Shanbhag SS, J Chodosh and HN Saeed. (2019). Sutureless amniotic membrane transplantation with cyanoacrylate glue for acute Stevens-Johnson syndrome/toxic epidermal necrolysis. Ocul Surf.

44. Kakabadze Z, D Chakhunashvili, K Gogilashvili, K Ediberidze, K Chakhunashvili, K Kalandarishvili and L Karalashvili. (2019). Bone Marrow Stem Cell and Decellularized Human Amniotic Membrane for the Treatment of Nonhealing Wound After Radiation Therapy. Exp Clin Transplant 17:92–98.

45. Vacz G, A Cselenyak, Z Cserep, R Benko, E Kovacs, E Pankotai, A Lindenmair, S Wolbank, CM Schwarz, DB Horvathy, L Kiss, I Hornyak and Z Lacza. (2016). Effects of amniotic epithelial cell transplantation in endothelial injury. Interv Med Appl Sci 8:164–171.

46. Mendis AH, TJ Venaille and BW Robinson. (1990). Study of human epithelial cell detachment and damage: effects of proteases and oxidants. Immunol Cell Biol 68 (Pt 2):95–105.

47. Guillomot M, E Campion, A Prezelin, O Sandra, I Hue, D Le Bourhis, C Richard, FH Biase, C Rabel, R Wallace, H Lewin, JP Renard and H Jammes. (2014). Spatial and temporal changes of decorin, type I collagen and fibronectin expression in normal and clone bovine placenta. Placenta 35:737–47.

48. Canciello A, L Greco, V Russo and B Barboni. (2018). Amniotic Epithelial Cell Culture. Methods Mol Biol 1817:67–78.

49. Kurosawa H. (2007). Methods for inducing embryoid body formation: in vitro differentiation system of embryonic stem cells. J Biosci Bioeng 103:389–98.

50. Wernig M, A Meissner, R Foreman, T Brambrink, M Ku, K Hochedlinger, BE Bernstein and R Jaenisch. (2007). In vitro reprogramming of fibroblasts into a pluripotent ES-cell-like state. Nature 448:318–24.

51. Zhao B, W Zhang, Y Cun, J Li, Y Liu, J Gao, H Zhu, H Zhou, R Zhang and P Zheng. (2018). Mouse embryonic stem cells have increased capacity for replication fork restart driven by the specific Filia-Floped protein complex. Cell Res 28:69–89.

52. Hoveizi E and S Ebrahimi-Barough. (2019). Embryonic stem cells differentiated into neuron-like cells using SB431542 small molecule on nanofibrous PLA/CS/Wax scaffold. J Cell Physiol.

53. Lee HJ, SM Nam, SK Choi, KY Seo, HO Kim and S-H Chung. (2018). Comparative study of substrate free and amniotic membrane scaffolds for cultivation of limbal epithelial sheet. Scientific Reports 8:14628.

